# Identification of disulfidptosis-related-ferroptosis associated lncRNAs signature as a novel prognosis model for kidney renal clear cell carcinoma

**DOI:** 10.1101/2023.06.06.543974

**Authors:** Zihan Xu

## Abstract

**Background:** Clear cell renal cell carcinoma (ccRCC) represents 80% of all kidney cancers and has a poor prognosis. Newly discovered types of programmed cell death, ferroptosis and disulfidptosis, could have a direct impact on the outcome of KIRC cancer. Long non-coding RNAs (lncRNAs), which possess stable structures, can influence cancer prognosis and might be potential prognostic prediction factors for KIRC cancer. This study aims to investigate the correlation between disulfidptosis-related ferroptosis-related lncRNA and ccRCC in terms of immunity and prognosis.

**Methods:** Coexpression analysis was employed to identify disulfidptosis-related ferroptosis-related long non-coding RNAs (DRFRLs). Differential expression analysis of DRFRLs was conducted using the ‘limma’ package in R software, and the ‘ConsensusClusterPlus’ package was utilized to identify molecular subtypes. Prognostic DRFRLs were identified via univariate Cox analysis, and a prognostic model based on eight DRFRLs was constructed through Cox regression analysis and the least absolute shrinkage and selection operator (LASSO) algorithm. Kaplan-Meier (K-M) survival curve analysis and receiver operating characteristic (ROC) curve analysis were utilized to evaluate the prognostic power of this model. Additionally, differences in biological function were investigated using Gene Ontology (GO) and Kyoto Encyclopedia of Genes and Genomes (KEGG), while immunotherapy response was measured by utilizing tumor mutational burden (TMB) and tumor immune dysfunction and rejection (TIDE) scores. Single-cell analysis from the Tumor Immune Single Cell Center (TISCH) was employed to investigate cells with specific expression of the eight identified lncRNAs.

**Results:** Two clusters (A and B) of disulfidptosis-related ferroptosis-related long non-coding RNAs (DRFRLs) were identified. Survival analysis revealed that patients with subtype A had a higher probability of survival compared to those in subtype B, suggesting that subtype A predicts better survival. An eight-lncRNA signature was established through LASSO-Cox regression, and Kaplan-Meier curves validated the accuracy of prognostic features prediction (P < 0.001). This signature demonstrated excellent prognostic performance, with an area under the curve (AUC) of 0.762, 0.761, and 0.749 at 1, 3, and 5 years in the training set and 0.790, 0.739, and 0.726 in the testing set, respectively. In the single-cell dataset, LINC01534, FOXD2-AS1, AC002070.1, and AL158212.3 were found to be expressed, with FOXD2-AS1 and AC002070.1 specifically expressing in the KC tumor immune microenvironment.

**Conclusions:** The proposed signature of eight lncRNAs is a promising biomarker for predicting clinical outcomes and therapeutic responses in patients with ccRCC.

## Introduction

Renal cell carcinoma (RCC) is one of the most common types of urinary tract cancer worldwide, affecting approximately 430,000 individuals in 2020[1]. Over 100,000 patients with RCC die each year due to cancer progression[2]. Clear cell renal cell carcinoma (ccRCC), the most prevalent histological subtype of renal cell carcinoma, easily metastasizes and has a poor prognosis for patients[3]. The main treatment for ccRCC is surgery**Error! Reference source not found.**, but the course of the disease is often marked by uncertain prognoses due to tumor recurrence or metastasis. Roughly one-third of patients present with metastases at the time of diagnosis[5].

Unfortunately, surgery can be difficult to remove renal cell carcinoma metastases, and recurrence frequently occurs after nephrectomy. In addition, unlike other urologic tumors, ccRCC is insensitive to both radiotherapy and chemotherapy[6]. Although immunotherapy has made significant breakthroughs in the treatment of ccRCC, the effectiveness of the treatment varies from person to person[7]. Given the limitations of surgery, chemotherapy, radiotherapy, and immunotherapy in treating ccRCC, it is crucial to explore alternative diagnostic biomarkers and potential therapeutic targets. Additionally, developing a risk model that can be customized for individualized patient treatment and predict prognosis is important. Recent studies have demonstrated the association between ferroptosis and ccRCC.

Ferroptosis was first proposed by Dixon in 2012, and the molecular compounds erastin and Ras selective lethal 3 (RSL3) can induce ferroptosis through different mechanisms. Erastin inhibits solute carrier family 7 member A11 (SLC7A11), which prevents cystine import and limits glutathione (GSH) synthesis[8]. ccRCC has been found to be highly sensitive to ferroptosis. This is due to its high dependence on GSH, and inhibition of cystine import and subsequent GSH synthesis can induce ferroptosis, which selectively reduces the viability of ccRCC[9]. Additionally, ferroptosis plays an essential role in ccRCC immune infiltration[10]. Furthermore, ferroptosis is linked to another newly discovered form of programmed cell death known as disulfidoptosis. Disulfidoptosis is caused by insufficient glucose intake and excessive cystine intake[11]. Cells that express high levels of SLC7A11 can inhibit ferroptosis when glucose is limited by taking up cystine through SLC7A11. However, this process may also lead to the induction of disulfidoptosis[11]-[12].

There may be some relationship between ferroptosis and disulfidoptosis, and balancing the two could become a new treatment strategy to improve outcomes. Therefore, a thorough study of the relationship between ferroptosis and disulfidoptosis would be beneficial in predicting the prognosis of ccRCC.

LncRNAs are frequently utilized in constructing prognostic models for various types of tumors. LncRNAs, which are non-coding RNA molecules longer than 200 nucleotides[13], play a vital role in regulating the transcription and translation of metabolism-related genesIts effects are mainly involved in regulating the transcription and translation of metabolism-related genes[14]. During tumor progression, lncRNAs can function as either oncogenes or tumor suppressors, thereby controlling tumor proliferation, differentiation, invasion, and metastasis[15]. Although an increasing number of studies have developed predictive models of ferroptosis-related lncRNAs[16]–[17], the biological behavior and prognosis of disulfidoptosis-related ferroptosis-related lncRNAs in ccRCC has yet to be explored. Based on the above analyses, we screened disulfidoptosis-related-ferroptosis-related lncRNAs associated with the prognosis of KIRC. Furthermore, we analyzed the prognostic signature of these disulfidoptosis-related-ferroptosis-related lncRNAs in KIRC samples to further enhance the prognostic markers of KIRC and provide innovative insights for clinically targeted drug therapy.

## Methods

### Data source

Publicly available expression and corresponding clinical data of patients with KIRC from TCGA (https://portal.gdc.cancer.gov/) were downloaded from the TCGA database up to April 12, 2023. The RNA-Seq data of 614 samples, which comprised of 72 normal samples and 542 tumor samples, were obtained. Next, Ensembl IDs were converted to official gene symbols, and the data was log2 transformed. LncRNAs and protein-coding genes were screened using the Ensembl human genome browser GRCh38.

### Identification of disulfidptosis-related-ferroptosis-related LncRNAs

The list of ferroptosis-related genes, which contained 259 validated human FRGs, was downloaded from FerrDb (http://www.zhounan.org/ferrdb/index.html)[18](Zhou and Bao, 2020), and is listed in Table S1. The 23 disulfidptosis-related genes, listed in Table S2, were obtained from previous disulfidptosis-related publications[11]. A correlation analysis was conducted between the genes associated with disulfidptosis and ferroptosis using the limma package at |cor| > 0.6 and P < 0.05, as shown in Table S4. To further identify potential disulfidptosis-related ferroptosis-related lncRNAs, co-expression correlation analysis of lncRNAs and disulfidptosis-related ferroptosis-related gene expression profiles was performed using the limma package at |R^2^|>0.4 and p < 0.001) [19]. The 625 related lncRNAs are listed in Table S3(|cor| >0.6 and P<0.05).

### Analysis of Differential Expression and potential prognostic DRFRlncRNAs

The limma package [20] was utilized to screen the expression matrix of disulfidptosis-related-ferroptosis-related lncRNAs between KIRC samples and normal kidney samples. The criteria for DElncRNAs were defined as |log2(fold change)| > 1 and a false discovery rate (FDR) <0.05[21]. The 204 Differential Expression LncRNAs that met the criteria were listed in Table S5. These 204 Differential Expression LncRNAs were further filtered by Cox univariate analysis using the ‘survival’, ‘survminer’, and ‘limma’ R packages to identify potential prognostic DRFRDElncRNAs (p < 0.05). A total of 267 patients were randomly divided into training or validation cohorts at a 1:1 ratio. The 59 potential prognostic-related DRFRDElncRNAs are listed in Table S6.

### Consensus clustering analysis of potential prognostic related DRFRDElncRNAs

Consensus clustering analysis was conducted on the disulfidptosis-related ferroptosis-related DElncRNAs listed in Table S6. The ‘ConsensusClusterPlus’ package in R was utilized to perform unsupervised clustering analysis and classify KC patients into different molecular subtypes based on the expression of potential prognostic-related DRFRDElncRNAs. The following criteria were used: firstly, ensuring that the number of samples in each group was relatively consistent. Secondly, ensuring that the cumulative distribution function (CDF) curve rose gradually and smoothly. Thirdly, confirming that after clustering, the intra-group link was stronger, while the inter-group link was weaker.

### Correlation of molecular Subtypes and prognosis

To investigate the clinical significance of molecular subtypes, Kaplan-Meier plots were generated and log-rank tests were performed to assess the correlation between molecular subtypes and overall survival (OS) of KC patients.

### Construction of disulfidptosis-related ferroptosis-related LnCRNAs Prognostic Signature

Potential prognostic DRFRLncRNAs (p < 0.05) were identified by filtering the disulfidptosis-related ferroptosis-related DElncRNAs through Cox univariate analysis using the ‘survival’ R package. A total of 267 patients were randomly divided into training or validation cohorts at a 1:1 ratio. The least absolute shrinkage and selection operator (LASSO)-Cox regression analysis was then applied to these prognostic candidates. Finally, an eight-LncRNA optimal prognostic model was established by selecting the penalty parameter λ that correlated with the minimum 10-fold cross-validation. The formula for DRFRLncRNAs-related prognostic risk scores for each patient was

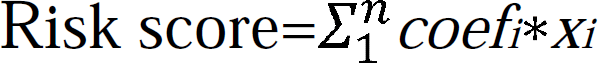

The expression of each lncRNA and its corresponding coefficient were represented by *xi* and *coefi*, respectively. Based on the median value of the risk score, patients in the training cohort were classified into low-risk and high-risk groups. The Kaplan-Meier curve was produced using the ‘survminer’ R package, with the log-rank test used to compare overall survival (OS) between the high/low-risk group. A receiver operating characteristic curve (ROC)[22]was generated to assess the predictive accuracy of the signature via the ‘timeROC’ R package. To evaluate the model’s feasibility, the risk score was computed in the validation cohort using the same formula as the training cohort, and then the same validation method was applied as described above.

### Development of a nomogram scoring system

The clinicopathological features and DRFRLncRNAs -related prognostic risk score was incorporated to develop a nomogram using the ‘rms’ package, based on patients’ survival. In the nomogram scoring system, a variable, such as gender, age, TNM stage, and CRG Risk score, was matched with a score, and the total score was obtained by adding the scores across all variables of each sample. The subsequent calibration graph of the nomogram scoring system was performed to examine the predictive value between the predicted 1-, 3-, and 5-year survival rates and the virtual outcomes.

### Functional Enrichment Analysis

Genes that exhibited differential expression between the high-risk and low-risk groups were identified using the ‘edgeR’[23]R package,with criteria set at |log2(fold change)| > 1 and FDR < 0.05. These genes were functionally annotated based on the Gene Ontology (GO) and the Kyoto Encyclopedia of Genes and Genomes (KEGG) using the ‘clusterProfiler’ R package[24], with the adjusted p-value threshold set at <0.05.

### Tumor Immune Dysfunction and Exclusion score

The TCGA website was used to obtain the somatic mutation file. And the original mutation annotation format (MAF) was divided into two groups according to the risk score. The aforementioned analyses were conducted using the maftools R package. The potential immune checkpoint blockade (ICB) response was evaluated using the tumor immune dysfunction and exclusion (TIDE) algorithm, and TIDE data for kidney clear cell carcinoma (KIRC) were downloaded from http://tide.dfci.harvard.edu/. The TIDE algorithm is a recent development that employs TIDE scores to accurately predict the efficacy of immunotherapy drugs administered to patients[25]. Jiang et al. found that TIDE scores have been shown to represent the efficacy of immune drugs (anti-PD1, anti-CTLA4) in melanoma patients, with higher TIDE scores being associated with better outcomes[25]Finally, we used the pRRophetic R package[26]to calculate the semi_inhibitory concentration (IC50) values of chemotherapeutic drugs.

### Exploring of Hub LncRNAs by Single-Cell Sequencing

We used the Tumor Immune Single Cell Hub (TISCH) project, developed by Sun et al[27], to examine the expression pattern of the prognostic model associated with the 8 LncRNAs at the single-cell level of the KC microenvironment. TISCH is a comprehensive database that combines single-cell transcriptomic profiles of 76 high-quality tumor datasets from 27 cancer types. The analyzed single-cell cohort comprised GSE111360, GSE145281_aPDL1, and GSE139555.

## Results

### Prognosis-Related lncRNAs With Coexpression of disulfidptosis-related ferroptosis

Seventy genes associated with disulfidptosis and ferroptosis have been detected. (Table S4). By performing co-expression analysis with stringent criteria (|Pearson R| > 0.6 and p < 0.05), we identified a set of 625 LncRNAs that were significantly co-expressed with the 70 genes of interest (Figure 2A). Further analysis revealed 204 differentially expressed long non-coding RNAs among these 625 candidates, based on the criteria of |logFC|>1 and FDR<0.05 when comparing tumor and normal tissues (Figure 2B and C). By utilizing univariate Cox analysis (p < 0.05), we selected several differentially expressed lncRNAs associated with disulfidptosis-related ferroptosis-related prognostic-related lncRNAs, including AC108673.3, FOXD2-AS1, AC002070.1, AF196972.1, AC124854.1, AP001267.3, LINC01583, AC008555.1, LINC01801, AL109741.1, AC007743.1, AC009093.1, AC007861.1, AC004816.1, AC010809.2, AC000050.2, AC068051.1, AL359715.3, AP001625.2, AC003984.1, AC147067.1, AC112220.2, AC017099.2, SUCLG2-AS1, AL158212.3, LINC01534, C3orf36, AL354811.1, AL589745.1, AC007365.1, AC004837.2, SIAH2-AS1, RAP2C-AS1, AC009486.1, AC027601.2, AL162377.1, AC007637.1, AC010333.2, AF111167.2, AL732509.1, TNFRSF10A-AS1, AC007406.3, AC007376.2, IL10RB-DT, AC145098.1, LINC01704, AL049840.5, HID1-AS1, LINC01624, AL162171.1, AC004918.3, LINC01415, CACTIN-AS1, LINC01738, LINC02521, AL355803.1, AL353648.1, AC027702.1, and AL133227.1 (Figure 2D).

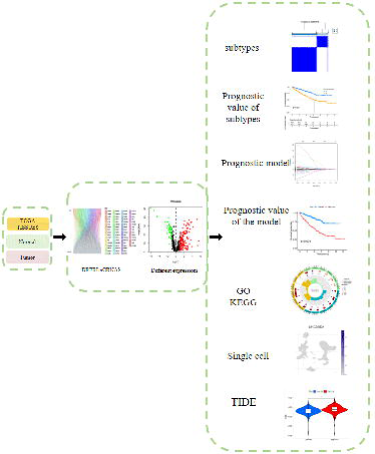

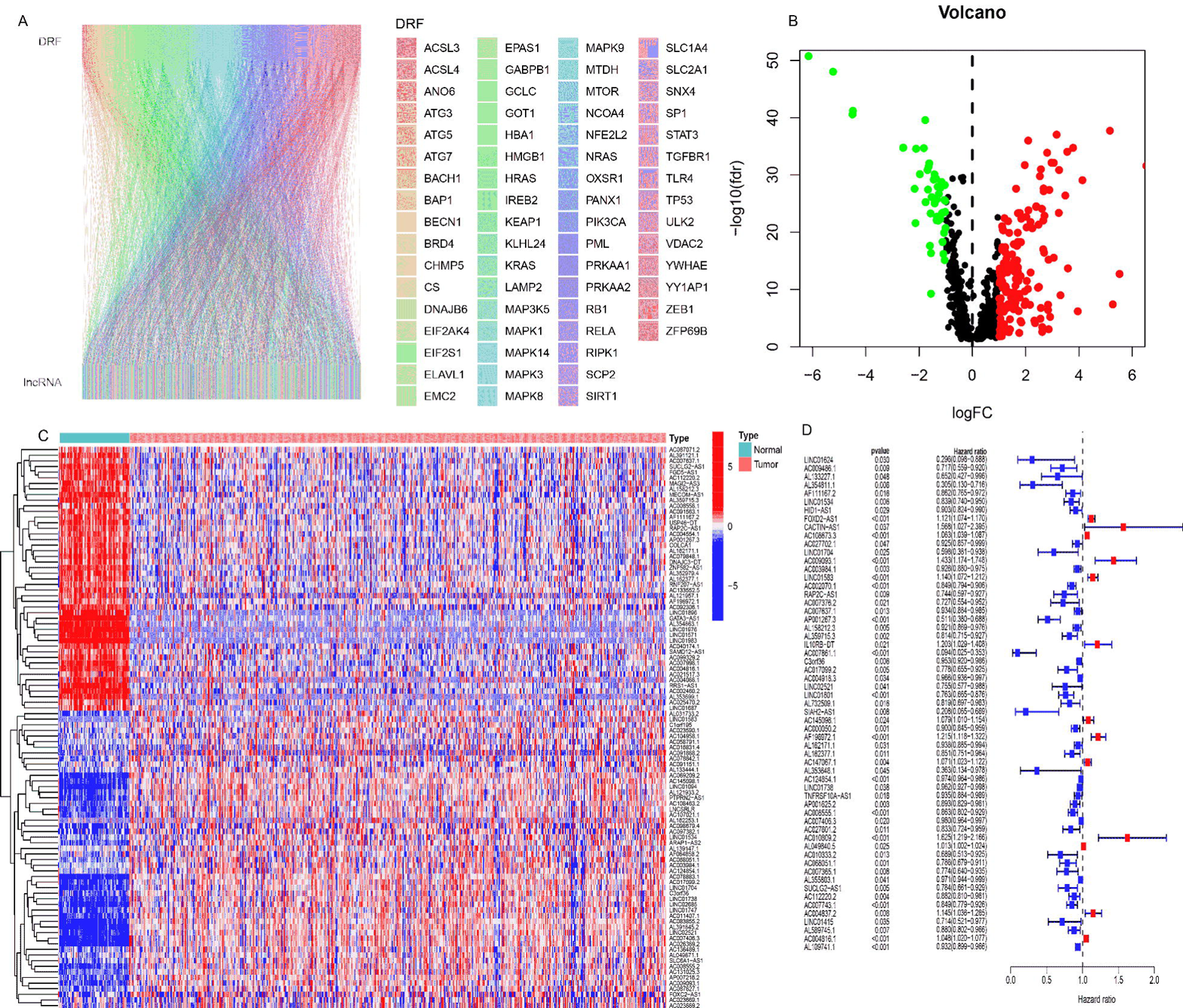

### Identification of disulfidptosis-related ferroptosis molecular subtypes

A disulfidptosis-related ferroptosis-related prognostic-related lncRNAs network was carried out to comprehensively demonstrate the association among DRFLncRNAs and their prognostic value in KC patients (Figure 3A). This network revealed the presence of prevalent and complex interactions among DRFLncRNAs. Utilizing a consensus clustering algorithm, we classified KC patients into two groups based on the expression profiles of DRFLncRNAs. Our analysis indicated that k=2 was the optimal selection for classifying patients into two distinct molecular subtypes, A (n=186) and B (n=428) (Figure 3B, Figures S1A–J; Table S7). UMAP and tSNE analysis confirmed significant differences in the in the disulfidptosis-related transcription profiles between subtype A and B (Figure 3C, Figures S2A). Furthermore, our survival analysis revealed that KC patients with subtype A had a higher probability of survival than those with subtype B (log-rank test, p < 0.001; Figure 3D).

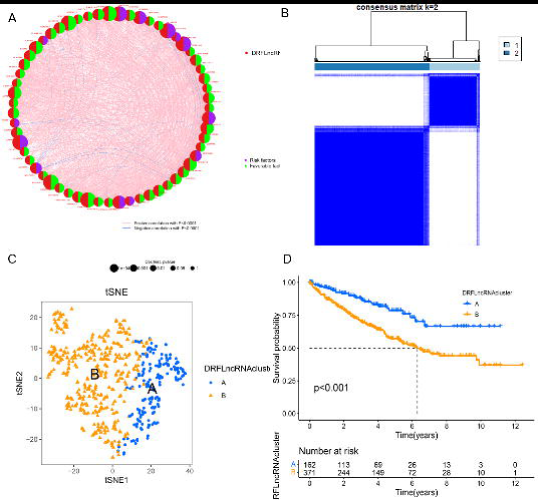

### Construction and Validation of a DRFLncRNAs Prognostic Model

To evaluate the prognostic value of DRFLncRNAs, we randomly divided KIRC samples from the TCGA database into two groups: a training group and a validation group. Using the optimal penalty parameter (λ) for the LASSO model from the 59 prognostic DRFLncRNA lesions mentioned previously, we constructed a prognostic risk evaluation model using only 8 DRFLncRNAs in the training group. The cvfit and lambda curve are shown in Figures 4A and B. Each KIRC patient in the TCGA database was assigned a risk score utilizing the following formula in this model: Risk Score = LINC01534 * (-0.26501) + FOXD2-AS1 * 0.11155 + CACTIN-AS1 * 1.4379 + AC002070.1 * (-0.16625) + AL158212.3 * (-0.074935) + AL162171.1 * 0.077080 + AC124854.1 * (-0.0156464) + AC010333.2 * (-0.33588) (Note: the name of each lncRNA indicates its expression level in the TCGA database). We performed Cox univariate and multivariate regression analyses to evaluate the independent predictive potential of this signature. Firstly, Cox univariate regression analysis revealed that this signature’s risk score was associated with OS rates in KIRC patients (p < 0.001; Figure 4C). Furthermore, multivariate Cox regression analysis indicated that this 8-DRFLncRNAs risk signature, along with stage M and age, could serve as an independent prognostic factor for predicting the OS rates of KIRC patients in the TCGA database (p < 0.001; Figure 4D). A predictive nomogram calculated the likelihood of survival for these patients by adding up the scores identified on the points scale for multiple related factors. The 1-, 3- and 5-year OS rates could be accurately predicted when compared to those of the ideal predictive model (Figures 4E, F).

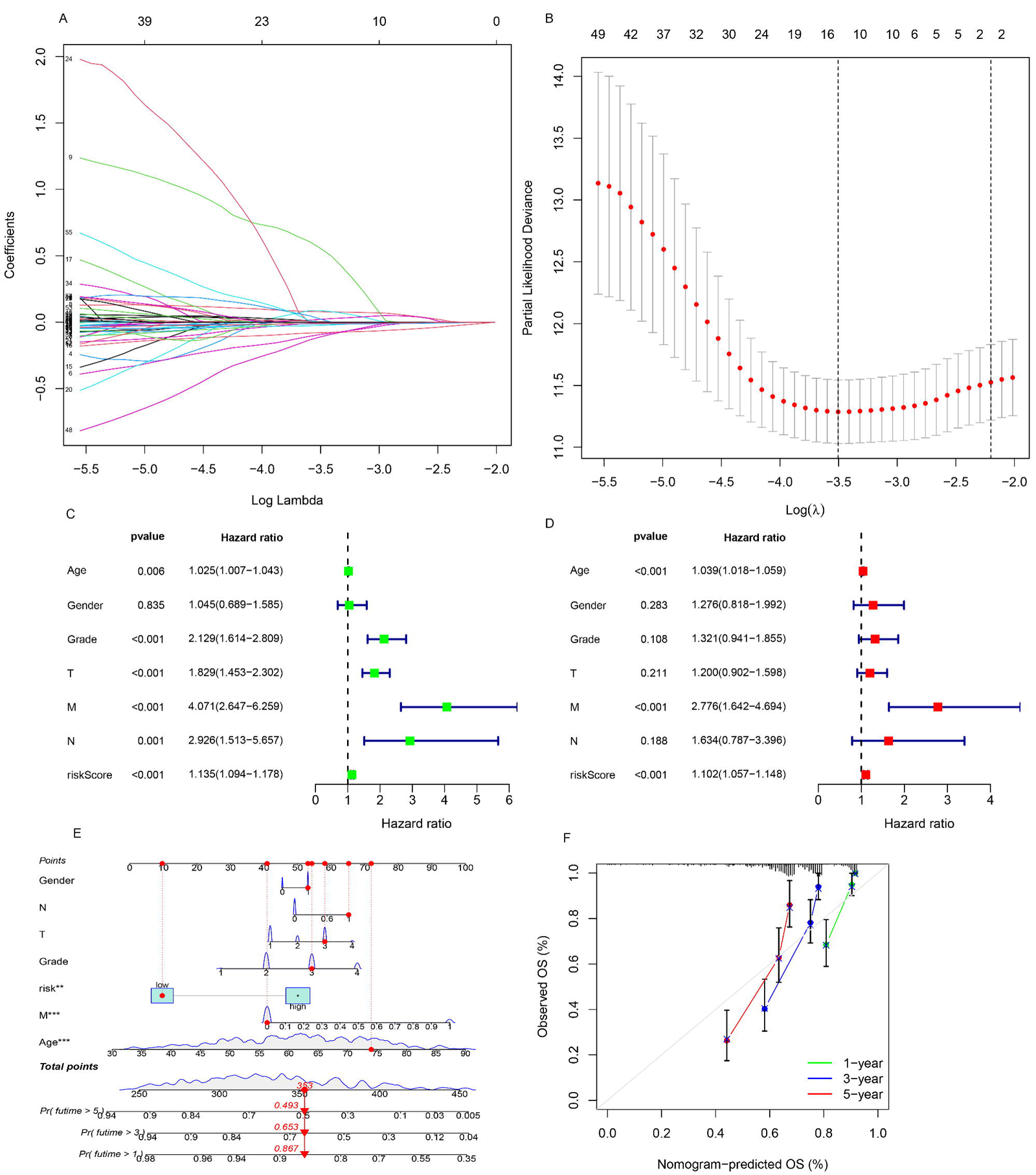

Subsequently, we evaluated the prognostic value of the 8-DRFLncRNAs model. The samples in the training group were classified into high-risk and low-risk groups according to the median value of the risk scores. The distribution of the risk scores and the distribution of the overall survival (OS) status were visualized to demonstrate that the two risk groups had a reasonable distribution (Figure 5A). Kaplan-Meier survival analysis was performed to show that patients with KIRC in the high-risk group had worse OS rates than those in the low-risk group (Figure 5D). Additionally, a time-dependent receiver operating characteristic (ROC) curve was generated for the training group. The areas under the curve (AUCs) were maintained at 0.762, 0.761, and 0.749 for the 1-year, 3-year, and 5-year points respectively (Figure 5G). To further evaluate the predictive efficacy of this 8-lncRNA signature, the distribution figures, Kaplan-Meier survival analysis, and ROC analysis were double validated in both the test group and the overall group. The samples in the test group also had a reasonable distribution in the risk groups (Figures 5B, E, H, K), with AUCs maintained at more than 0.72. In the overall group, the samples in the risk groups were also reasonably distributed (Figure 5C, F, I), with AUCs maintained at more than 0.73. These results suggest that individuals in the high-risk group may have higher mortality rates than those in the low-risk group.

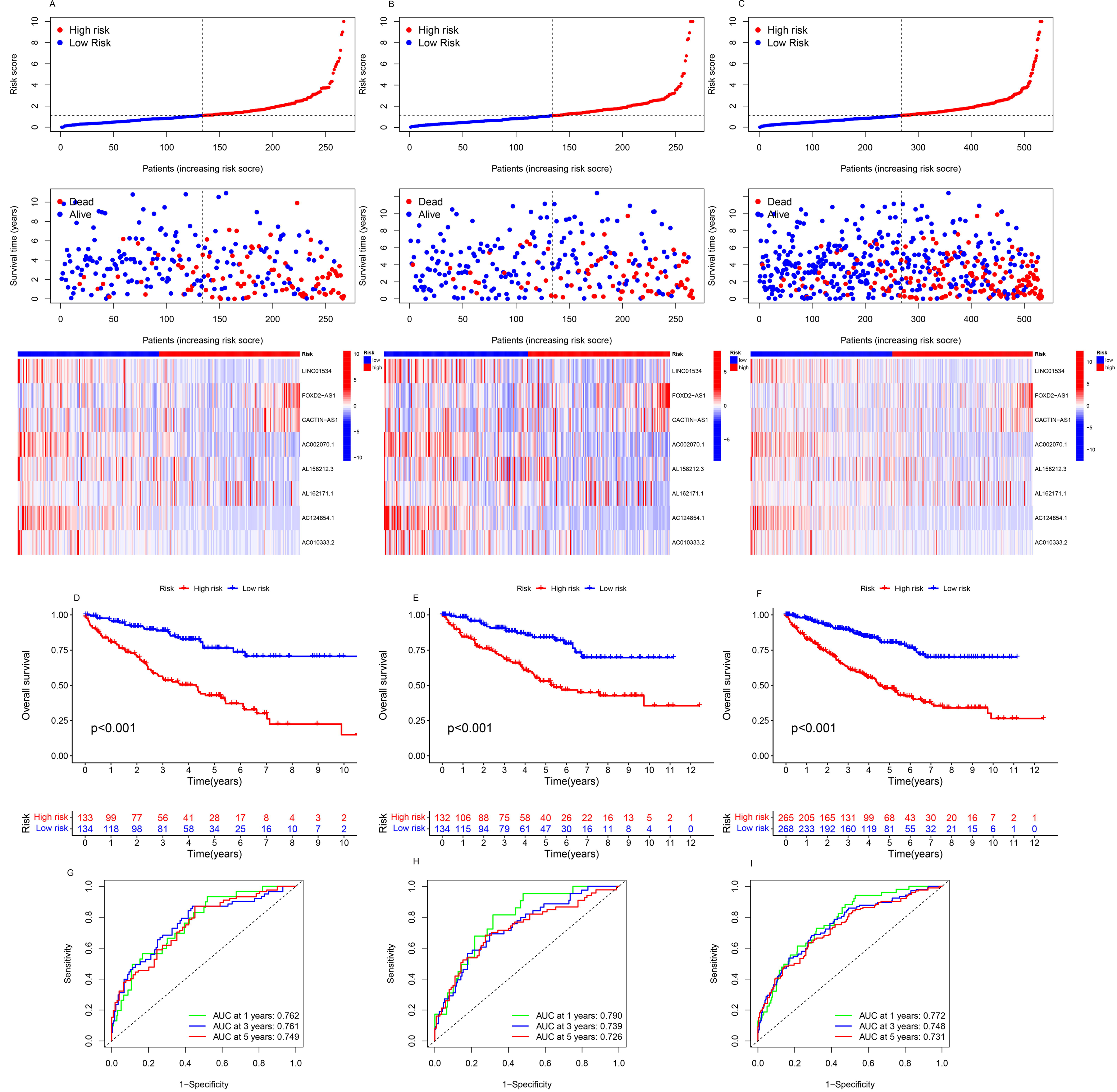

### Discovery of molecule function and pathways by gene ontology and kyoto encyclopedia of genes and genomes analysis

To explore the biological functions characterizing differentially expressed genes (DEGs) between high-risk and low-risk groups, we employed a log2|FC| > 1 and FDR < 0.05 cutoff to identify DEGs. Subsequently, GO enrichment analysis and KEGG pathway analysis were performed (p < 0.05). The GO analysis revealed significant enrichments in biological processes (BP), molecular functions (MF), and cell components (CC), as presented in Figure 6A table S8. Notably, immune-related functions were significantly enriched, particularly regarding antigen binding, immunoglobulin receptor binding, immunoglobulin complex, and B cell receptor signaling pathway. Similarly, the KEGG analysis identified three main pathways: protein digestion and absorption, fat digestion and absorption, and the TGF-beta signaling pathway, which is an immune-related pathway (Figure 6B table S9). Taken together, these results suggest that the risk score of the 8-lncRNAs signature is primarily associated with tumor immunity, tumor metastasis, and biological metabolism in KIRC.

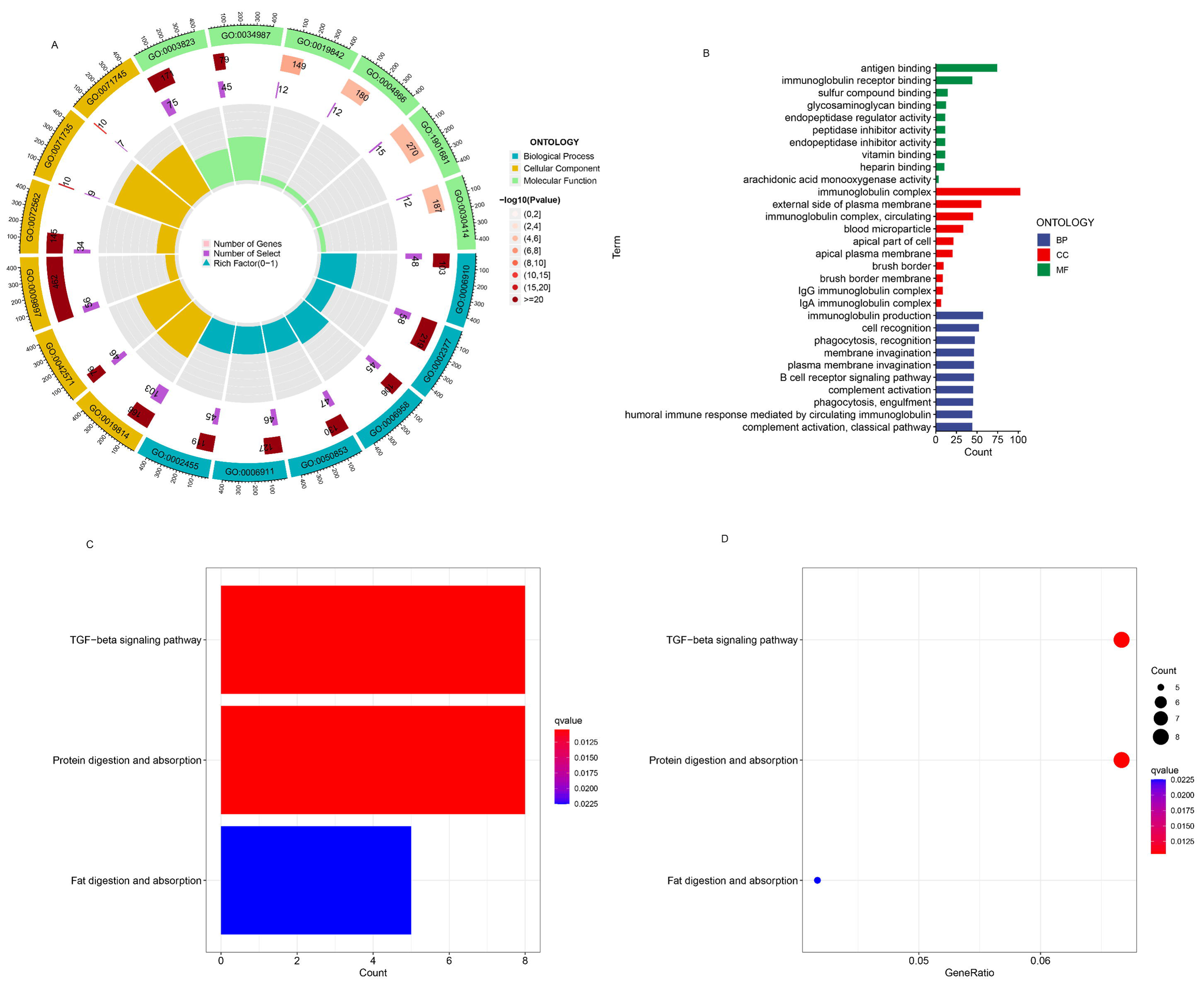

### TMB, TIDE, and Therapeutic Drug Sensitivity

In the context of immune checkpoint suppression therapy being a focus of cancer treatment, we investigated the ability of risk models based on disulfidptosis-related-ferroptosis-associated lcnRNAs to predict tumor-associated immunity, as presented in Figure 7. We obtained somatic mutation data from the TGCA database and analyzed changes in somatic mutations between high- and low-risk groups. The five most highly mutated genes in the high-risk group were VHL, PBRM1, TTN, SETD2, and BAP1 (Figure 7A and B). Among these genes, VHL, PBRM1, SETD2, and BAP1 were the most frequently mutated genes in ccRCC. However, there was no significant difference in TMB detected between the two groups (Figure 7C). Interestingly, TIDE scores were dramatically higher in the high-risk group compared to the low-risk group (Figure 7D). Analysis of 13 immune-associated pathways revealed significant differences in type I and II IFN response, HLA, checkpoint, co-stimulation, cytolytic activity, pro-inflammation, APC, CCR, and paraneoplastic inflammation between the high- and low-risk groups (Figure 7E). We further investigated drug sensitivity and found significant differences in IC50 values between the low- and high-risk groups for multiple drugs. Supplementary Figures S4 and S3 show the drugs sensitive to the high-risk group and low-risk group, respectively. It is worth noting that we chose two drugs that were identified as effective in the high-risk group (Figure 8A, B) and two drugs that were effective in the low-risk group (Figure 8C, D).

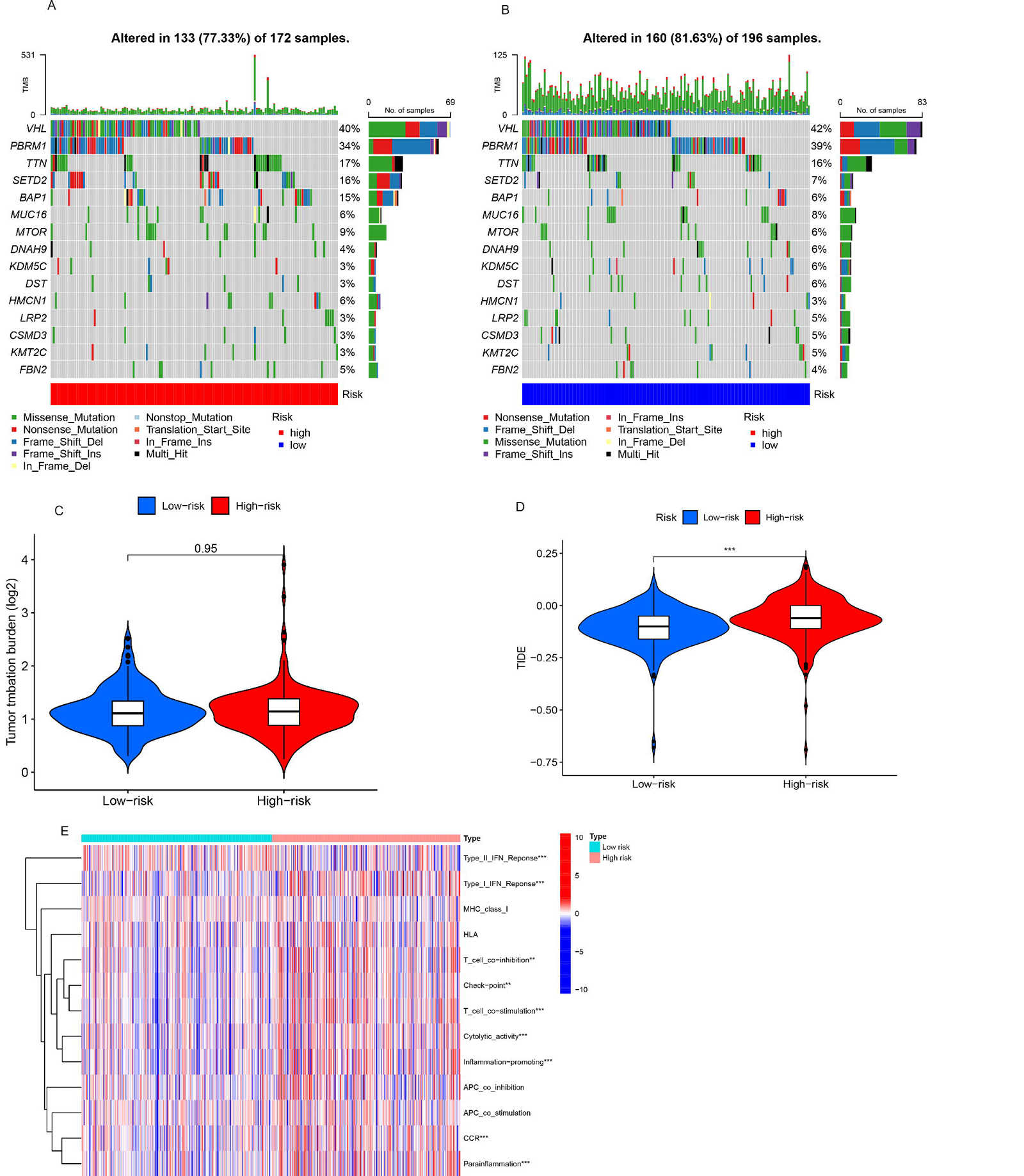

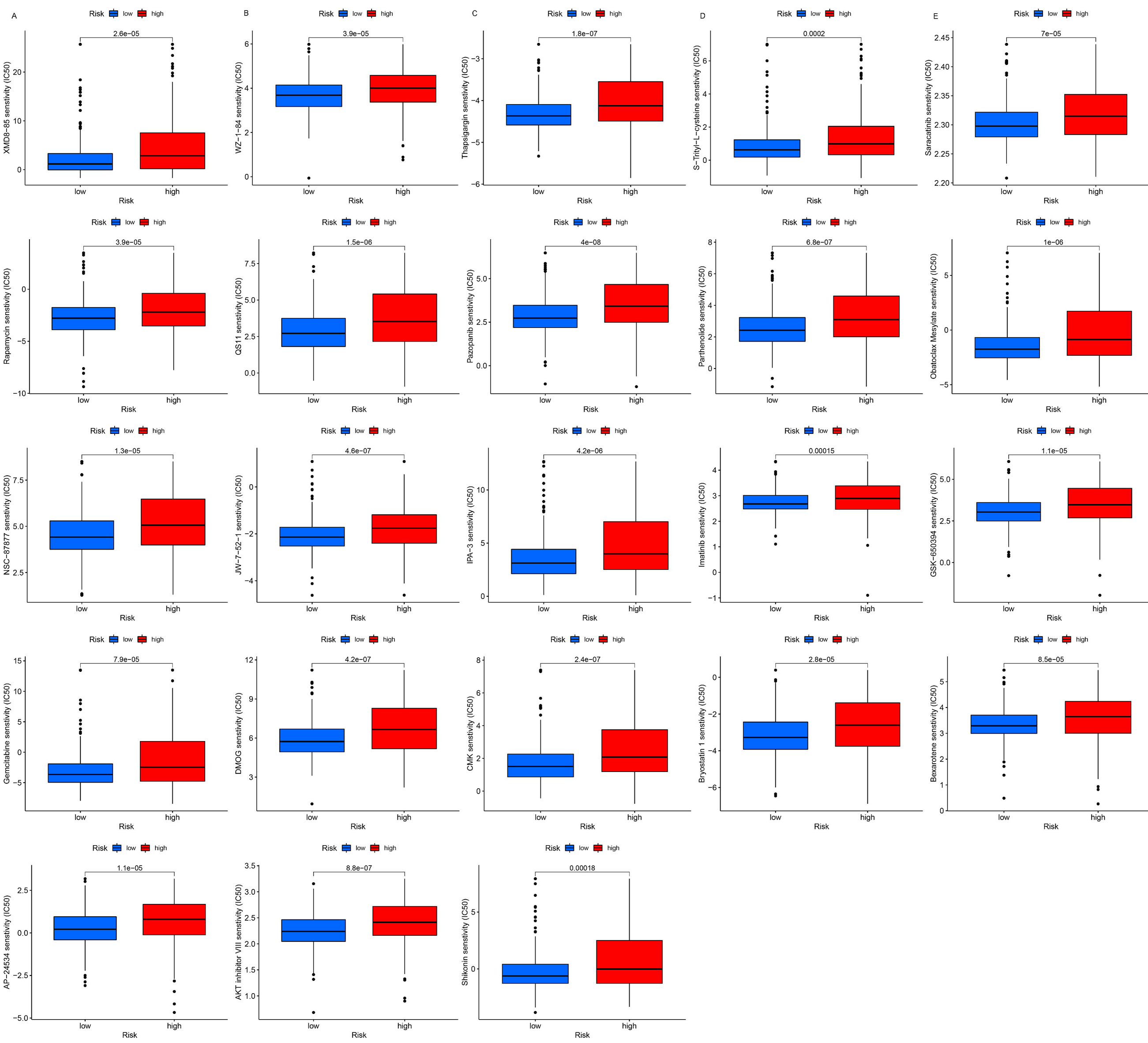

### Single-cell RNA analysis

To determine the cellular location of the 8 lncRNAs mentioned above, we performed single-cell RNA analysis using the Tumor Immune Single Cell Hub (TISCH) project. The GSE111360 dataset was initially divided into 24 cell clusters, as shown in Figure 9A. Based on cell markers, a total of 12 types of cell subpopulations were identified, as presented in Figure 9B. Furthermore, datasets GSE139555 and GSE145281_aPDL1 were also utilized in Figure S5. Among these datasets, four of the eight lncRNAs were identified: LINC01534, FOXD2-AS1, AC002070.1, and AL158212.3 (Figure 9C, D, E, F). FOXD2-AS1 and AC002070.1 were found in both GSE111360 and the post-immunotherapy dataset GSE145281_aPDL1. Notably, FOXD2-AS1 was expressed in most immune cells (Figure 9D), suggesting that it may play an important role in the development of ccRCC.

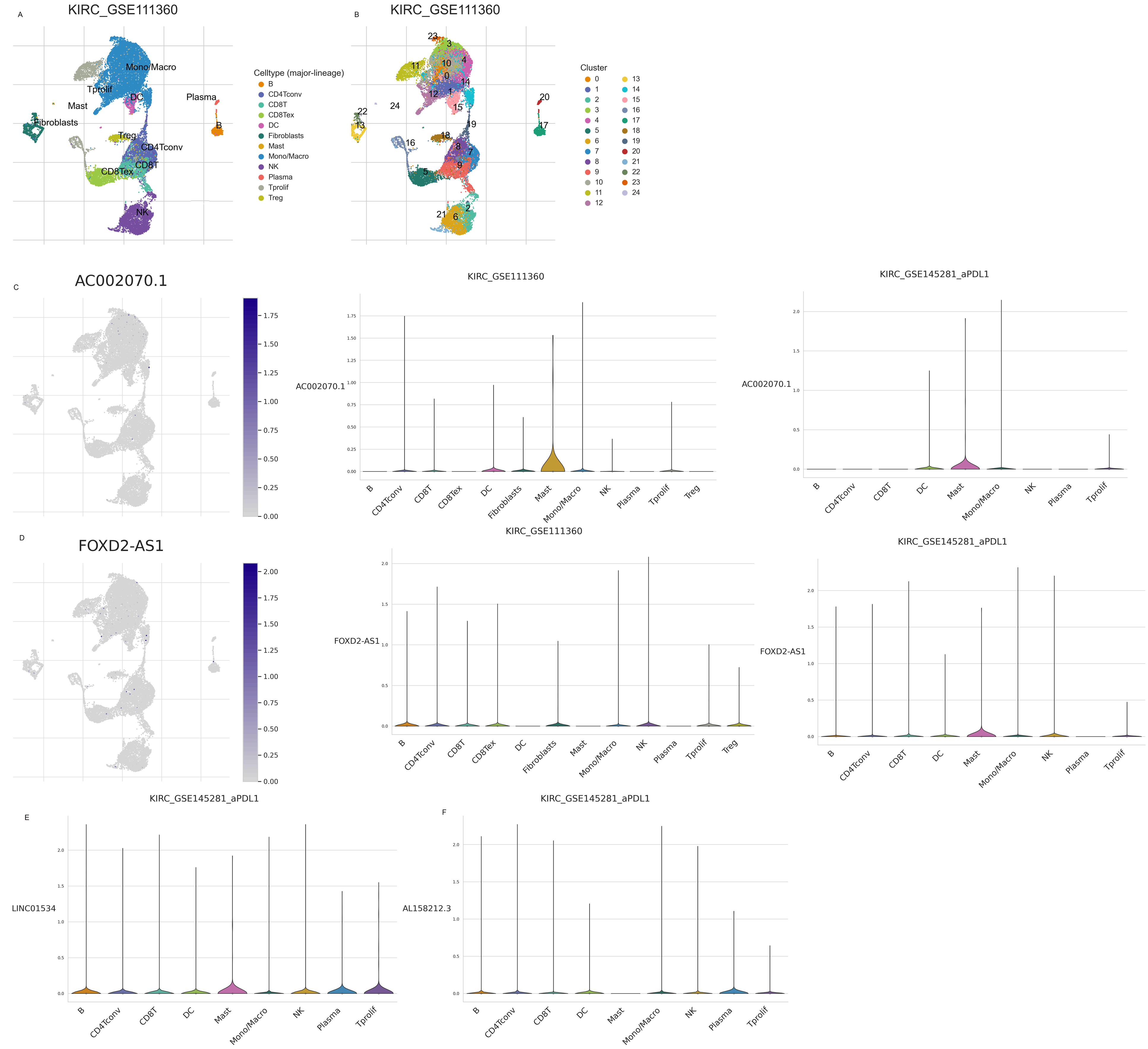

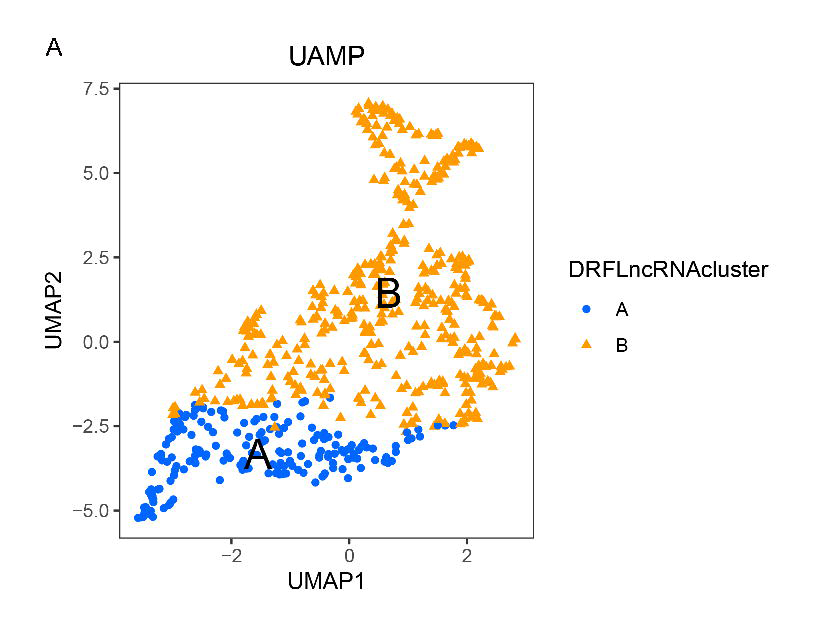

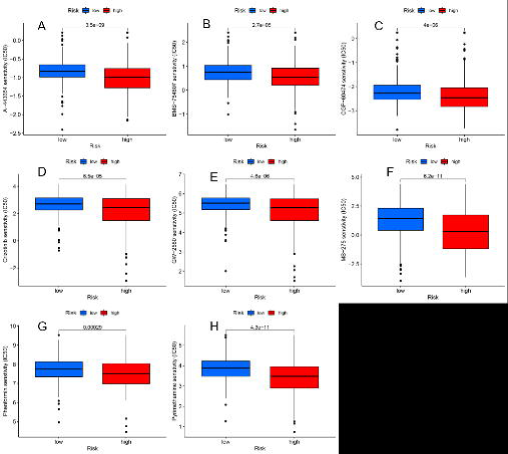

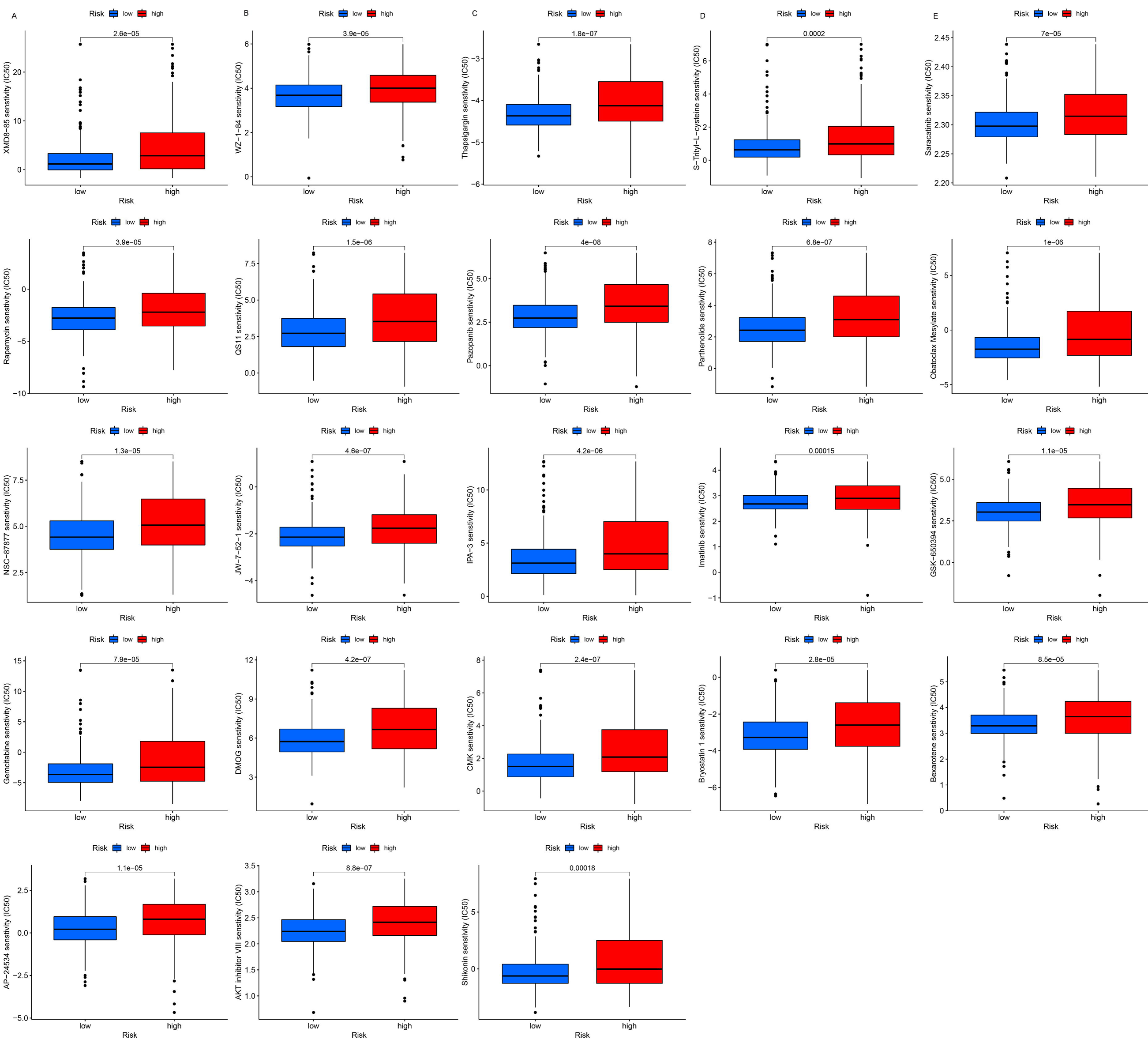

## Discussion

Long non-coding RNAs (lncRNAs) are known to play a crucial role in ccRCC, regulating genes and signaling pathways via various mechanisms, including miRNA-mediated pathways[28]. Several regulatory axes of lncRNAs have been identified that control the invasion and metastasis of ccRCC, such as CDKN2B-AS1/miR-141/CCND1[29] and RP11-436H11.5/miR-335-5p/BCL-W[30]. Researchers have also investigated ferroptosis-related lncRNAs in ccRCC to identify new biomarkers and establish prognostic models[31]-[32], while others have aimed to inhibit ccRCC migration and growth by inducing ferroptosis through specific targets [33].

Clear cell renal cell carcinoma (ccRCC) is a metabolic dysregulation disease [34]. Recently, cell death through disulfidptosis has been associated with metabolic processes, and a therapeutic strategy for bone metabolic diseases has been proposed by inducing disulfidptosis. Specifically, the study suggests that TXNRD1 occurs by selectively inducing disulfidptosis in osteoclast precursors to prevent osteoclast differentiation. As there is a balance between disulfidptosis and ferroptosis [11]-[12], it may be possible to identify lncRNAs that play a balancing role between disulfidptosis and ferroptosis and investigate their regulatory roles in ccRCC. By finding a corresponding target drug to break the balance between disulfidptosis and ferroptosis, we can promote ferroptosis or disulfidptosis in tumor cells to suppress the progression and metastasis of ccRCC.

In our study, we aimed to investigate the relationship between lncRNAs associated with disulfidptosis-related ferroptosis and ccRCC. Using correlation analysis, we identified 70 ferroptosis genes associated with disulfidptosis (cor>0.6, P<0.05). Co-expression analysis revealed 625 lncRNAs associated with these 70 genes and some of these lncRNAs may play a balancing role in disulfidptosis and ferroptosis. We then searched for possible ccRCC biomarkers from this list of 625 lncRNAs. LncRNAs that were differentially expressed in tumor and normal tissues and associated with prognosis were selected to construct a prognostic model, resulting in an 8-lncRNA prognostic prediction model for KIRC cancer. Additionally, we examined the expression of these eight lncRNAs in different cells using single-cell data. Of note, FOXD2-AS1 and AC002070.1 were found in both GSE111360 and the post-immunotherapy dataset, with FOXD2-AS1 expressed in most immune cells.

Previous research has linked FOXD2-AS1 to cuproptosis[35]-[36] and shown that it is highly expressed in ccRCC cell lines[35]. FOXD2-AS1 is aberrantly expressed in various cancers and has been linked to cancer progression[37] via targeting Akt/E2F1[38], P53[39], the microRNA-98-5p/CPEB4 axis [40] and miR-25-3p/Sema4C [41]. Our study has identified FOXD2-AS1 as a long non-coding RNA that is associated with disulfidptosis and ferroptosis. Additionally, previous studies have demonstrated the correlation between FOXD2-AS1 and cuproptosis, as well as the overexpression of FOXD2-AS1 in renal clear cell carcinoma.

Excessive copper has been suggested to increase iron toxicity and the development of oxidative stress, which may be related to ferroptosis[42]. Some studies have suggested that CuO may promote apoptosis and cell cytotoxicity modulated by reactive oxygen species (ROS) [43]-[44]. According to the aforementioned evidence, it is suggested that ferroptosis, which is a type of cell death induced by the accumulation of ROS[45] may potentially interact with cuproptosis and copper-iron interactions in diverse physiological and pathological processes, including cancer progression.

Studies have shown that there may be a balance between ferroptosis and disulfidptosis. Therefore, we speculate that FOXD2-AS1 is able to balance disulfidptosis and ferroptosis through the interaction of cuproptosis and ferroptosis. In addition, 5 out of the 8 lncRNAs play important roles in cancer. For example, AL162171.1 is one of the costimulatory-related long non-coding RNAs in ccRCC[46]. AC124854.1 is a ferroptosis-related lncRNAs which is consistent with our findings [47]. One of the eight LnCRNAs we identified, LINC01534, was also found in the single-cell dataset and is expressed in tumor cells. In colorectal cancer tissues, LINC01534 is upregulated and plays a significant role in maintaining cancer stemness[48]. AL158212.3 is a M6-A related lncRNA in glioma [49], while CACTIN-AS1, AC002070.1, and AC010333.2 were first proposed in our study and had not been previously identified.

Predicting the drug sensitivity promoted improving drug selectivity and increasing the success rate of therapy [50]. Surprisingly, the high-risk group of patients was more susceptible to Cyclopamine, Bleomycin, Bexarotene, DMOG, and other drugs. In an earlier study, cyclopamine was well tolerated by mice[51]. In glioblastoma, Cyclopamine acts as a suppressant of carcinogenesis[52]. This may provide novel therapeutic strategies in KIRC patients.

In our study, we aimed to investigate the correlation between ferroptosis and disulfidptosis associated lncRNAs with ccRCC, and to identify biomarkers to create a prognostic model. However, there are still several issues that need to be addressed. Firstly, our research lacks the necessary experimental validation related to lncRNA expression and other relevant experiments. Furthermore, compared to protein-coding genes, lncRNAs are generally less conserved between species, which highlights the need for more prospective studies and basic research to further refine the relevant details of this study.

## Conclusions

In summary, we have identified long non-coding RNAs related to disulfidptosis and ferroptosis, and constructed a novel and accurate prognostic model. This signature may provide potential targets for enhancing immunotherapy in patients with clear cell renal cell carcinoma.

## Data availability statement

The original contributions presented in the study are included in the article/Supplementary Material. Further inquiries can be directed to the corresponding authors.

